# Gene editing in bacteria via SpCas9/sgRNA ribonucleoprotein complexes

**DOI:** 10.1101/2021.06.21.449263

**Authors:** Rubén Dario Arroyo-Olarte, Ricardo Bravo Rodríguez, Edgar Morales-Ríos

**Affiliations:** Departamento de Bioquímica, Centro de Investigación y Estudios Avanzados, Instituto Politécnico Nacional, Ciudad de México, México

**Keywords:** CRISPR-Cas, ribonucleoprotein, gene-editing, GFP, BFP

## Abstract

Gene editing has been revolutionized by CRISPR (Clustered Regularly Interspaced Short Palindromic Repeats)-Cas technology in a variety of organisms. In bacteria, however, CRISPR-Cas still holds many caveats such as high toxicity of Cas9 and off-target editing effects. In this work we develop a system for the incorporation of Cas9/single guide RNA ribonucleoprotein complexes in bacteria and their successful application for gene editing via homologous recombination. Targeting of a green fluorescent protein (GFP) reporter allows for easy verification of gene-edition via conversion to blue-fluorescent protein (BFP), mediated by the well characterized 196T > C (Tyr66His) mutation.

## 1. Introduction

Feasibility of the CRISPR-Cas system among prokaryotes varies greatly depending on several factors, e.g. Cas proteins cytotoxicity, AT genome content and available genetic transformation methods. In this regard, evaluation of loss of fluorescence in GFP-expressing bacteria serves as a straightforward way to assess CRISPR-Cas activity *in vivo* in different species. This has been applied in *E. coli* with a 2-plasmid system, one encoding for Cas9 and GFP-specific sgRNA expression, and another for GFP expression. GFP-plasmid loss varied between 80 % to 98 % of colonies depending on the sgRNA sequence [1]. The system can also be used to assess the efficiency of CRISPR-Cas-mediated gene editing. It has been shown that the Tyr66-His mutant (encoded by the single base substitution 196T>C) shifts wild-type GFP absorption and emission towards the blue spectrum, thus creating blue fluorescent protein (BFP) (Fig. 1) [2]. The loss of the hydrogen-bond provided by tyrosin side-chain hydroxyl group, also generates a slightly more relaxed tertiary structure in BFP (Fig. 1). This amino acid change is involved as well in the higher stability of BFP fluorescence at acidic pH, but lower at alkaline pH compared to GFP [3].

**Figure 1.**
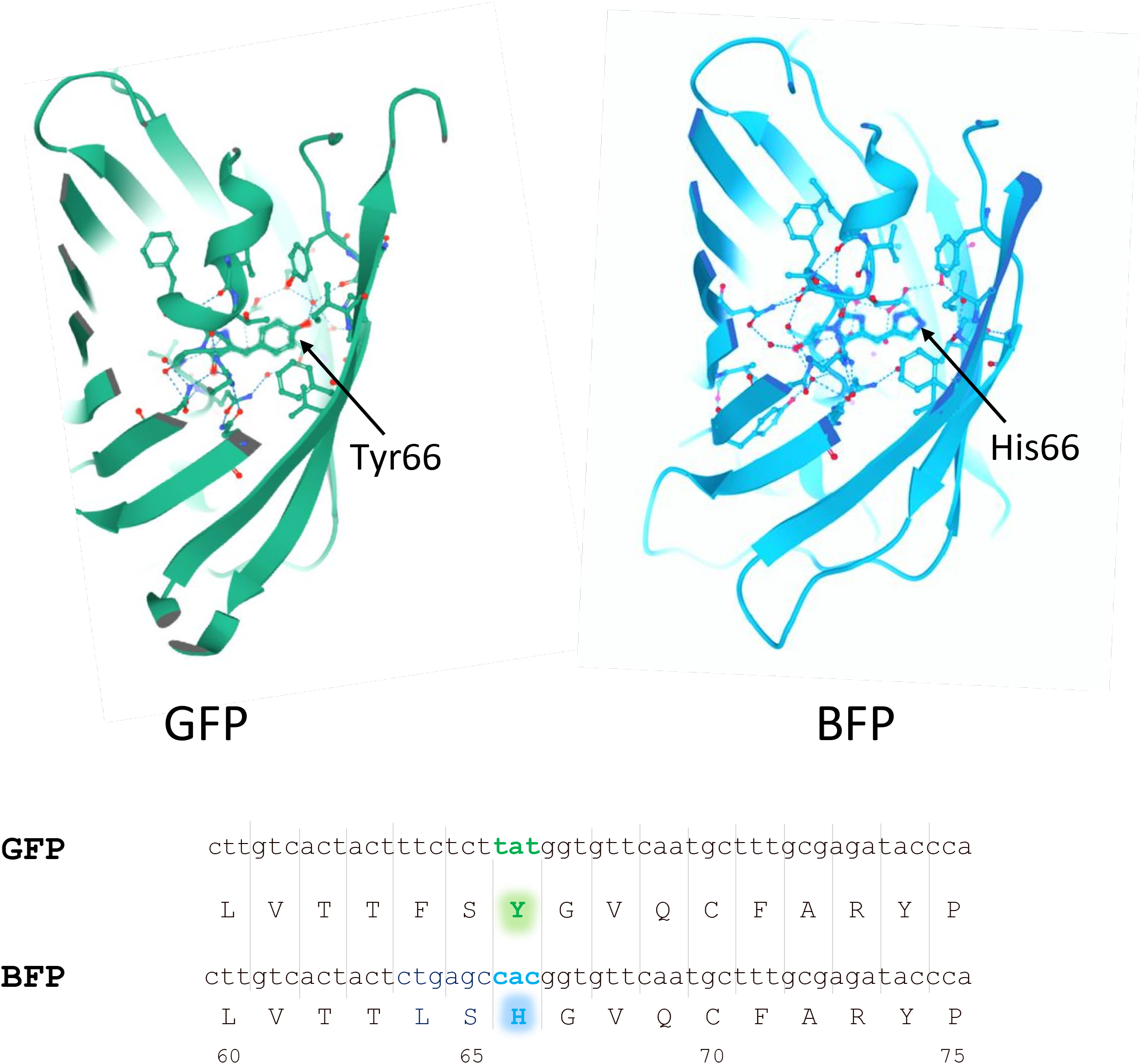
Converting the green fluorescent protein to the blue fluorescent protein. Crystallographic structures of the GFB and BFP shows the amino acid residues close to the tyrosine 66 in GFP and the histidine 66 in BFP. The Tyr66-His mutation (encoded by a single nucleotide substitution at 196T>C) has been shown to alter wild-type GFP uptake and blue spectrum luminescence, resulting in the blue fluorescent protein (BFP).

A GFP to BFP conversion assay has recently been applied to evaluate a plasmid-based CRISPR/Cas9 system in *Methylococcus capsulatus* [4]. Transformation with Cas9/single guide (sg)RNA ribonucleoprotein (RNP) complexes carries several advantages. The main advantage of this approach is that it does not rely on the host transcription and translation machinery. Besides, the RNP complex is usually degraded shortly after transfection, avoiding the toxic effects of a continuous Cas9 expression (Fig 4). It also does not require cloning. Therefore, there is no restriction in the selection of sgRNAs that may target a cloning strain genome. It also presents a more concise streamline than the plasmid methods, as no plasmid curing is required. In this work we applied the RNP approach in a GFP to BFP conversion assay [5] and show successful gene editing in *Escherichia coli*, and potentially in other prokaryotic organisms, by homologous recombination with an exogenous DNA repair template.

## 2. Materials

### 2.1. Plasmids, PCR, sgRNA *in vitro* transcription and nuclease assays

1. pET-NLS-Cas9-6xhis and pAKgfp1 are requested from Addgene (https://www.addgene.org), with ID no. 62934 and 14076, respectively. The pET-NLS-Cas9-6xhis plasmid is used to express recombinant *Sp*Cas9 for affinity purification, while pAKgfp1 plasmid is used to express GFP in *Escherichia coli*.
2. DNA oligonucleotides are requested to be used as homology-directed repair (HDR) template, for DNA template synthesis by overhang PCR for sgRNA *in vitro* transcription (IVT), and for Cas9 substrate PCR to be used in nuclease assays (Table 1):
3. TranscriptAid T7 high yield transcription kit (Thermo Fischer Scientific).
4. Common molecular biology reagents, e.g., nuclease-free water, ethyl alcohol, isopropyl alcohol, agarose, proteinase K, RNAseZAP™ (Sigma-Aldrich), Taq DNA polymerase, DNA purification spin-column system, ethidium bromide/SYBR green.
5. 10x Cas9 reaction buffer: 100 mM NaCl, 50 mM Tris-HCl, 10 mM MgCl_2_, 100 μg/ml BSA, pH 7.9.

**Table 1.**
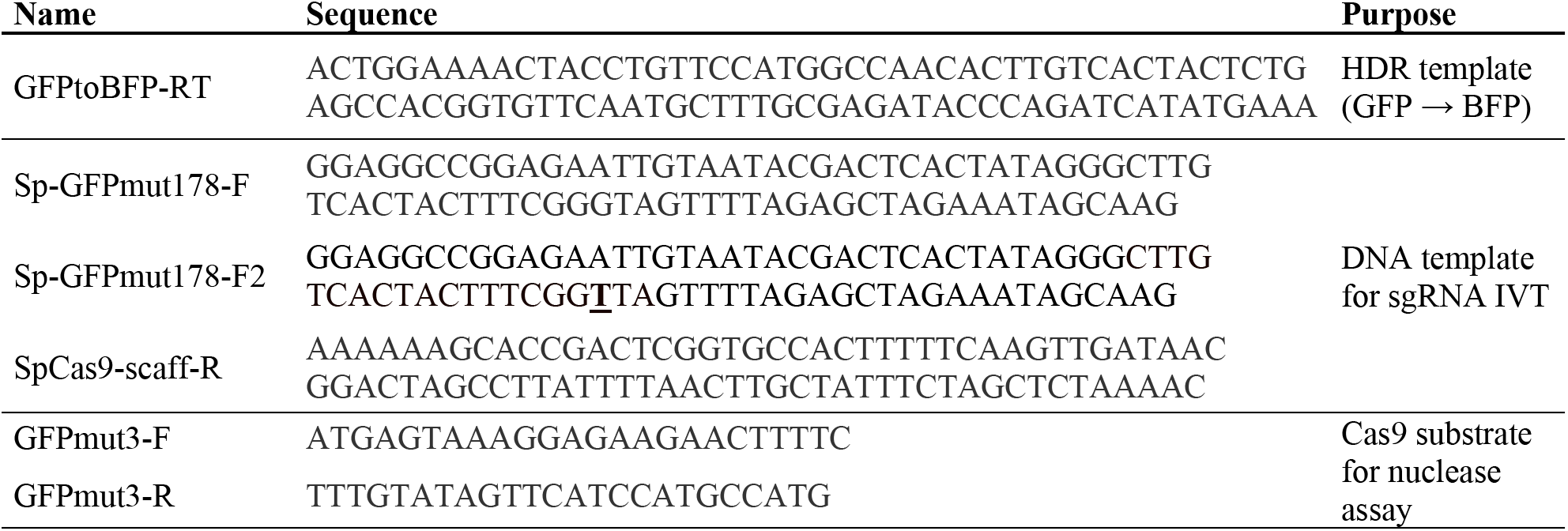
Oligonucleotides used in this study

### 2.2. *Escherichia coli* strains and culture

1. Competent bacterial cells: *Escherichia coli* strain soluBL21™ (Genlantis) for recombinant SpCas9-6xhis expression and purification, and *E. coli* strain TOP10™ (Thermo Fischer) for GFP expression and gene editing.
2. Luria-Bertani agar plates supplemented with ampicillin (Sigma-Aldrich).
3. 2xYT medium: 16 g/L tryptone, 10 g/L yeast extract, 5.0 g/L NaCl.
4. Isopropyl β-galactoside (IPTG, Sigma-Aldrich).
5. Swinging bucket centrifuge and microcentrifuge.
6. Temperature-controlled shaker and thermoblock.

### 2.3. *Escherichia coli* electroporation

1. 0.1 cm-gap sterile electroporation cuvettes (Biorad).
2. Gene Pulser X-Cell electroporation system (Biorad).
3. Phosphate-buffered saline (PBS) supplemented with 10 % glycerol.
4. SOC medium: 0.5% (w/v) yeast extract, 2% (w/v) tryptone, 10 mM NaCl, 2.5 mM KCl, 20 mM MgSO_4_, 20 mM glucose.

### 2.4. Recombinant SpCas9-6xhis purification

1. Ultrasonic homogenizer VC-500 (Cole-Palmer).
2. ÄKTA pure 25L purification system (Cytiva).
3. 5 ml Nickel-nitrilotriacetic acid (Ni-NTA) HisTrapTM column (Cytiva).
4. Gel filtration Superdex 200 increase column (Cytiva).
5. Amicon 100 KDa centrifugal filter units (Sigma-Aldrich).
6. Lysis buffer 10x: 300 mM Tris, 2 M NaCl, 20 mM MgCl_2_, pH 7.5.
7. Buffer A: lysis buffer 1x, 5 mM imidazole, 2 mM benzamidine, 0.8 mM phenylmethylsulfonyl fluoride (PMSF), 3 mM β-mercaptoethanol, pH 7.0.
8. Buffer B: lysis buffer 1x, 500 mM imidazole, 2 mM benzamidine, 0.8 mM PMSF, 3 mM, β-mercaptoethanol, pH 7.0.
9. Sodium dodecyl sulphate (SDS), polyacrylamide, electrophoresis chamber.
10. Cas9 storage buffer: 50 mM 4-(2-hydroxyethyl)-1-piperazineethanesulfonic acid (HEPES), 300 mM NaCl, 0.5 mM β-mercaptoethanol, 10% glycerol, 2 mM benzamidine, 0.8 mM PMSF.
11. Coomassie Brilliant Blue G-250 for Bradford protein assay.
12. UV-vis spectrophotometer and 1 cm-gap cuvettes.

## 3. Methods

### 3.1. Cell culture and SpCas9-6xhis expression

*Escherichia coli* strain soluBL21™ (Genlantis) was transformed by heat-shock (40-45 sec, 42 ⁰C) with plasmid pET-NLS-Cas9-6xhis (Addgene). Transformants were selected by culturing in Luria-Bertani agar plates supplemented with 150 μg/ml ampicillin (Sigma-Aldrich). Recombinant SpCas9-6xHis expression was induced by culturing in 2xYT medium (16 g/L tryptone, 10 g/L yeast extract, 5.0 g/L NaCl) supplemented with 1mM isopropyl β-galactoside (IPTG, Sigma-Aldrich), for 16-18 h at 18 ⁰C. Cells were centrifuged (4,000 xg, 30 min), and pellets were stored at −70 ⁰C.

### 3.2. Ni-NTA affinity and size-exclusion chromatography

Cell pellets were lysed by sonication (50 %, 10 min: 30 s pulse, 5 s rest). Lysate was centrifuged (30,000 xg, 30 min) to remove cell debris. Cleared lysate was filtered (0.2 μm Milipore) and affinity chromatography was performed on an ÄKTA pure 25L purification system (Cytiva). Sample was applied onto a 5 ml Ni-NTA HisTrap™ column (Cytiva) at a constant flow rate of 1.5 ml/min. Next, unbound proteins were washed out at a flow rate of 3 ml/min. A gradient solution containing imidazole from 5 mM to 500 mM was then applied to the column to elute bound proteins into 5 ml fractions. Real-time UV spectrophotometrically (260 nm) allowed to visualize the elution of different protein peaks.

Eluted affinity fractions were separated by SDS-PAGE and gels were stained with Comassie blue. Fractions showing at least 80 % purity of the expected SpCas9 protein band size (160 kDa) were merged and concentrated by centrifugation (3,200 xg) on a 100 kDa Amicon filter unit.

Size-exclusion chromatography was then performed on a gel filtration Superdex 200 column (Cytiva) to further purify SpCas9 and to remove small molecule contaminants from elution buffer (e.g., imidazole). Final SpCas9-6xHis was re-concentrated on a 100 kDa Amicon filter unit and dissolved in Cas9 storage buffer. Final protein concentration was determined by a Bradford protein assay on a linear range of BSA standards (0-20 μg/ml).

### 3.3. *In vitro* transcription of sgRNAs

First, the DNA template was generated by overhang PCR by mixing 1-unit Taq polymerase, 30 μl sgRNA-specific forward primer (10 μM Sp-GFPmut178-F/F2), 20 μl of reverse primer (10 μM SpCas9-scaff-R) and 1x reaction buffer. Forward and reverse oligonucleotides share a 25-bp overlapping sequence. The amplification protocol used will depend on the selected DNA polymerase, but a good approximation is as follows:

i. 94 °C for 2:00 minutes
ii. 94 °C for 30 seconds
iii. 55-58 °C for 30 seconds
iv. 68-72 °C for 25 seconds
v. Cycle steps ii to iv for 34 times
vi. 68-72 °C for 5 minutes

PCR product is cleaned up using a commercial spin-column based system and purified DNA template is then used for IVT according to manufacturer’s instructions. Briefly, 500-2000 ng DNA template are mixed with reaction buffer, 2 μl enzyme mix, 2 μl ATP, 2 μl CTP, 2 μl GTP and 2 μl UTP (75 μM each). Reaction is run at 37 °C for 5 h, and sgRNA is purified by ethanol precipitation. sgRNA purity is assessed by A_260_/A_280_ ratio (>1.8) and integrity is verified by agarose electrophoresis and UV visualization.

### 3.4. *In vitro* nuclease assay

The first step is the preparation of Cas9/gRNA RNP, by incubating the following mix at 25 °C for 15 min:

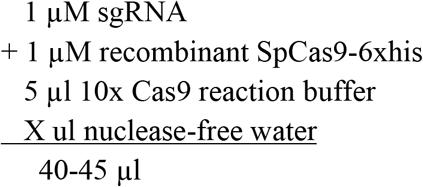

Thereafter 5-10 μl Cas9 substrate (GFPmut3 PCR product, 300 nM) are added, and reaction is incubated at 37 °C for 1h. Reaction is then stopped by adding 2 μl proteinase K (10 mg/ml) at room temperature for 20 min. Reaction products are then analyzed by agarose gel electrophoresis. Densitometric analysis of uncut vs. cut DNA bands observed under UV is performed with ImageJ software (NIH) to evaluate the reaction efficiency.

### 3.5. RNP electroporation

First the following reagents are mixed:

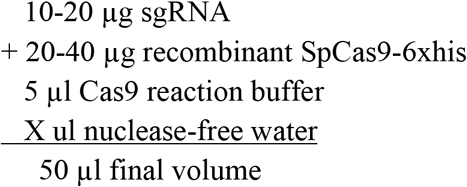

An initial incubation of 15 min at room temperature is performed to allow Cas9/gRNA RNP formation. Thereafter, 80-100 μg DNA repair oligonucleotide (GFP to BFP) are added and combined to 50 μl competent GFP-expressing *E. coli* TOP10 (in PBS/glycerol 10 %, 4 °C).

The cell suspension (100-120 μl) is then transferred to 0.1-cm cuvettes and electroporated in a Gene-PulserX (Bio-Rad) with the following parameters: 2 pulses, voltage 1.25 kV, capacitance 200 μF. 900 μl SOC medium are immediately added and transformed cells are allowed to recover for 2 h, 37 °C, 1,000 xg before plating in LB-agar plates supplemented with ampicillin (150 μg/ml).

## 4. Results

### 4.1. Purification of recombinant SpCas9

Protein samples at every stage of purification were collected and analyzed using SDS-PAGE (Fig. 2A-B). During the first step, Ni-NTA affinity chromatography, an elongated peak to the right was observed containing SpCas9 (Fig. 2A). Most contaminant proteins appeared in the fractions with less imidazole. As imidazole gradient increased, fractions started to show a 160 kDa band corresponding to SpCas9 as the dominant protein. During the second step, size-exclusion chromatography, there were three major peaks with elution volumes of 8.8, 13.6 and 16.3 ml (Fig. 2B). The first peak (8.8 ml) showed several bands including a 160 kDa protein in SDS-PAGE. However, this elution volume was too low for the expected molecular size of monomeric SpCas9 on the Superdex 200 increase 10/300 GL column (Figure S1) and was assumed to be comprised of soluble aggregates. Fractions corresponding to the second peak (13.6 ml) showed almost exclusively a 160 kDa band which also corresponds to the expected elution volume of similar-size proteins globular proteins purified by the same gel-filtration column (Table S1). The third peak (16.3 ml) showed a mixture of contaminant proteins of lower molecular size (Fig. 2B). There was also a fourth minor peak eluting at 20.6 ml that matched a conductance peak corresponding to the desalting of low-molecular weight imidazole (68 Da) carried on from the Ni-NTA affinity chromatography step. Finally, purified SpCas9-6xhis was concentrated to 3-5 μg/μl for further experiments.

**Figure 2.**
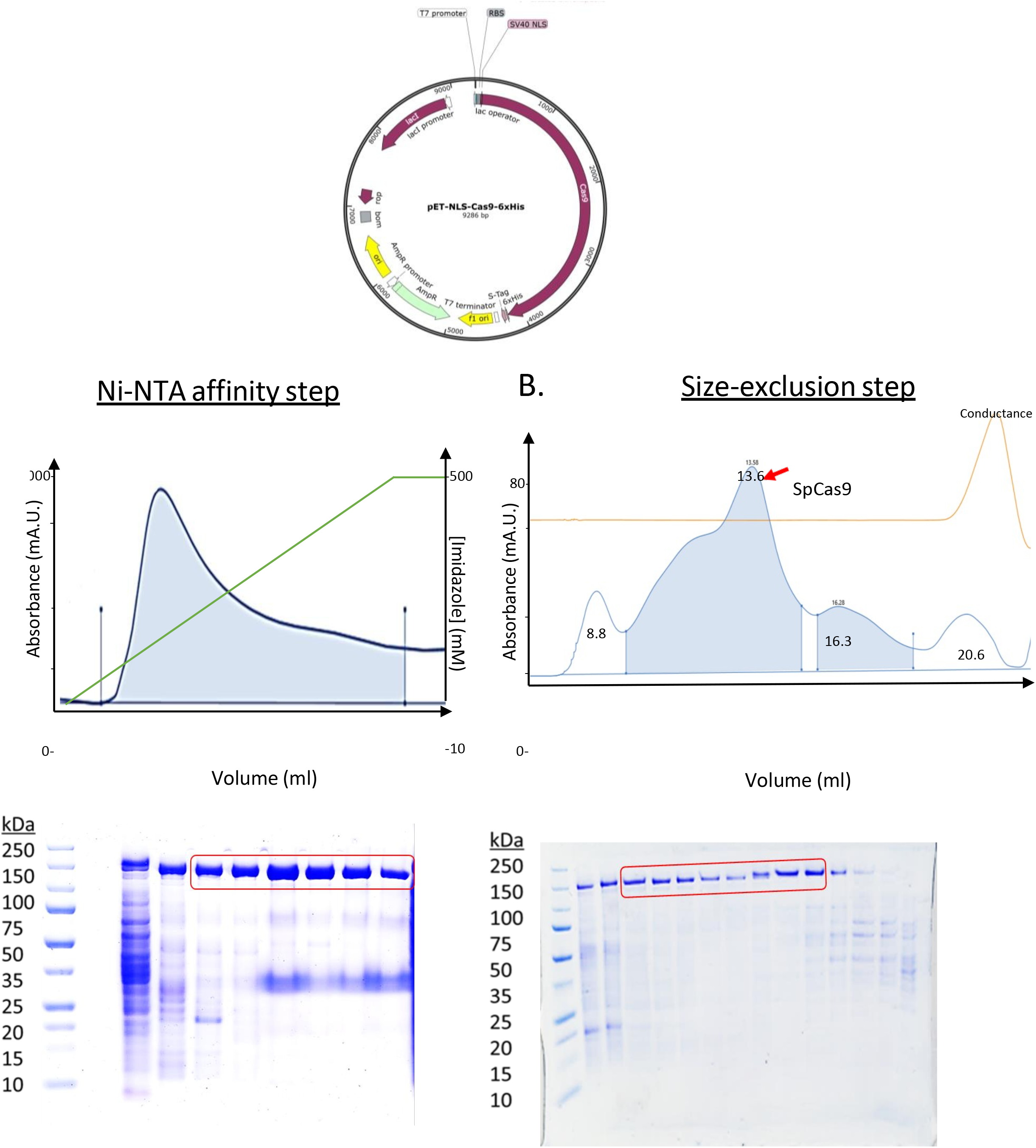
Purification of the recombinant Cas9 from Streptococcus pyogenes. During the first step, Ni-NTA affinity chromatography, an elongated peak to the right was observed containing SpCas9 (A) As the imidazole gradient increased, a band of 160 kDa appears on the fraction analysis by SDS-PAGE corresponding to SpCas9 as the major protein. In the second step, size exclusion chromatography, there were three main peaks with an elution volume of 8.8,13.6,16.3 ml (B). The first peak (8.8 ml) shows several bands containing the 160 kDa protein, the samples were collected and also analyzed by SDS-PAGE (Fig. 2A-B)The fraction corresponding to the second peak (13.6 ml) shows a band of 160 kDa. The third peak (16.3 ml) shows a mixture of contaminated low molecular size proteins (Fig. 2B). There is also a fourth minor peak eluting at 20.6 ml, consistent with the conductivity peak corresponding to the imidazole (68 Da).

### 4.2. *In vitro* nuclease assay shows high and specific activity of recombinant SpCas9

In order to analyze the efficacy of the purified SpCas9-6xHis as a potential gene-editing tool, nuclease assays were performed in combination or absence of a specific sgRNA. Our results show that our preparation has no background nuclease activity in absence of a sgRNA targeting the fluorochrome site of the GFP gene as DNA substrate (PCR product) (Fig.3).

**Figure 3.**
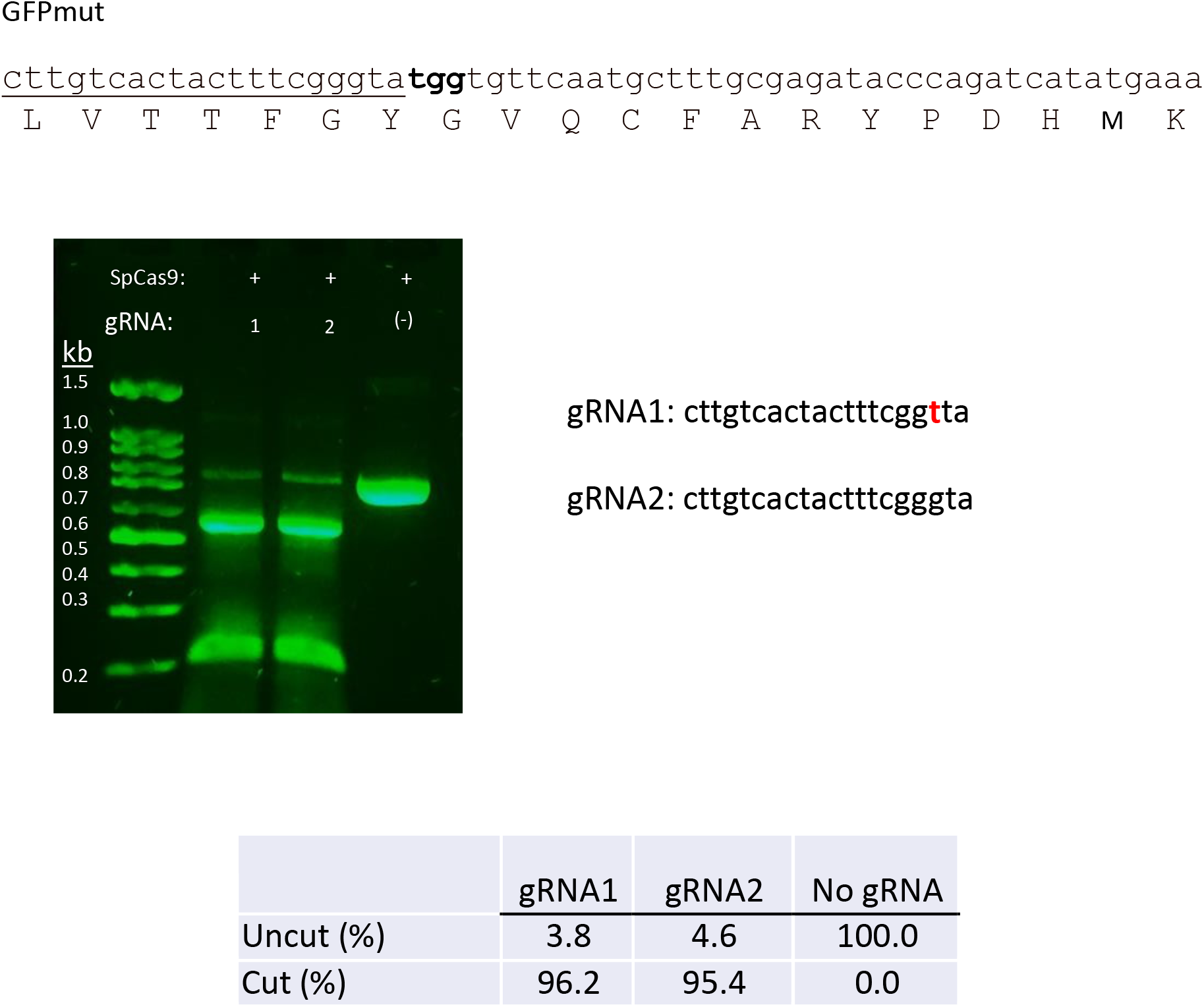
Characterization of the purified SpCas9. Our two-step purification method produced a high enzymatic activity SpCas9-6xHis. In the absence of sgRNA targeting the fluorochrome region of the GFP gene using a DNA substrate (PCR product), had no background nuclease activity. The RNP complex sgRNA and the DNA substrates were incubated and there was no difference in the efficiency of specific double-strand nuclease activity of both gRNAs (96.2% vs. 95.4%).

**Figure 4.**
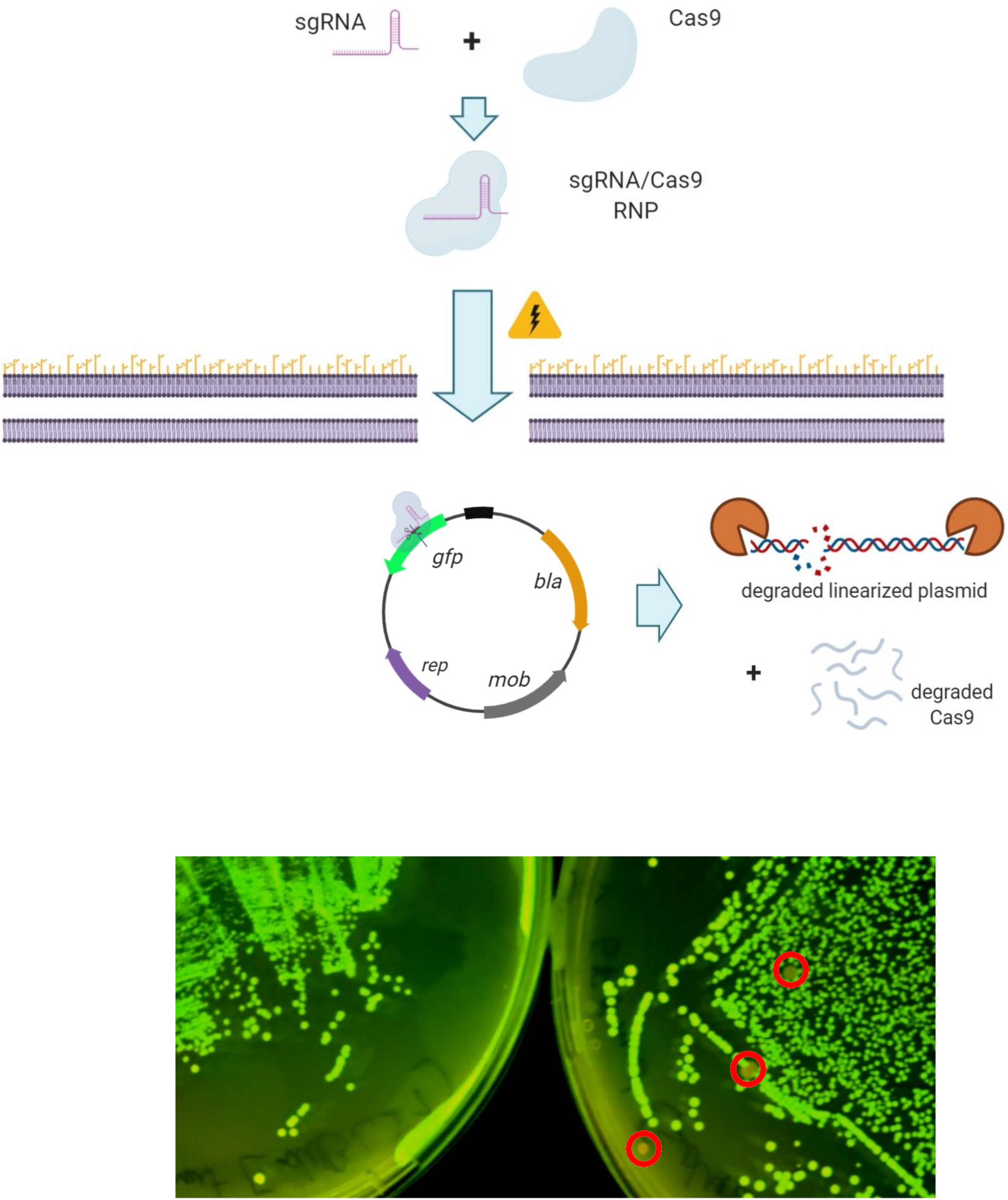
Electroporation of GFP-expressing bacteria with the RNP. The transformation Cas9 and sgRNA by electroporation resulted in loss of plasmid-expressed GFP fluorescence in *E. coli* colonies. A short DNA recombination template (30 bp homology arm) encoding the mutation responsible for the transition from GFP to BFP (Tyr66 → His) was also used to turn blue in RNP-transformed cells. These results are proof of the principle of use of RNP in bacteria.

However, when incubating the DNA substrate with RNP complexes of purified SpCas9-6xHis plus either a perfect-match or 1bp-mismatch (as close as 3 bp from PAM) sgRNA, there was no difference in the efficiency of formation of a specific double strand break (96.2 % vs 95.4 %) (Fig. 3).

### 4.3. Electroporation of RNPs and a DNA repair template induces GFP to BFP conversion

Transformation with the RNPs of a highly active Cas9 and sgRNA (Fig. 2C) by electroporation leads to colonies loss of plasmid-encoded GFP fluorescence in *E. coli* colonies (Fig. 4). Remarkably, when a short DNA recombination template (30 bp homology arms) encoding mutations responsible for the GFP to BFP transition (Tyr66→His) is also used, we could observe blue-fluorescent cells in RNP-transfected cultures (Fig. 5). Although the observed efficiency is low (<5 %), these results are a proof of principle for the use of RNPs in bacteria, which to our knowledge has not been reported so far in these organisms. As the transformation efficiency of a SpCas9/gRNA complex would expectedly be limited due to its massive size, recombineering efficiency must be maximized. However, CRISPR nuclease cytotoxicity would be minimized to achieve editing.

**Figure 5.**
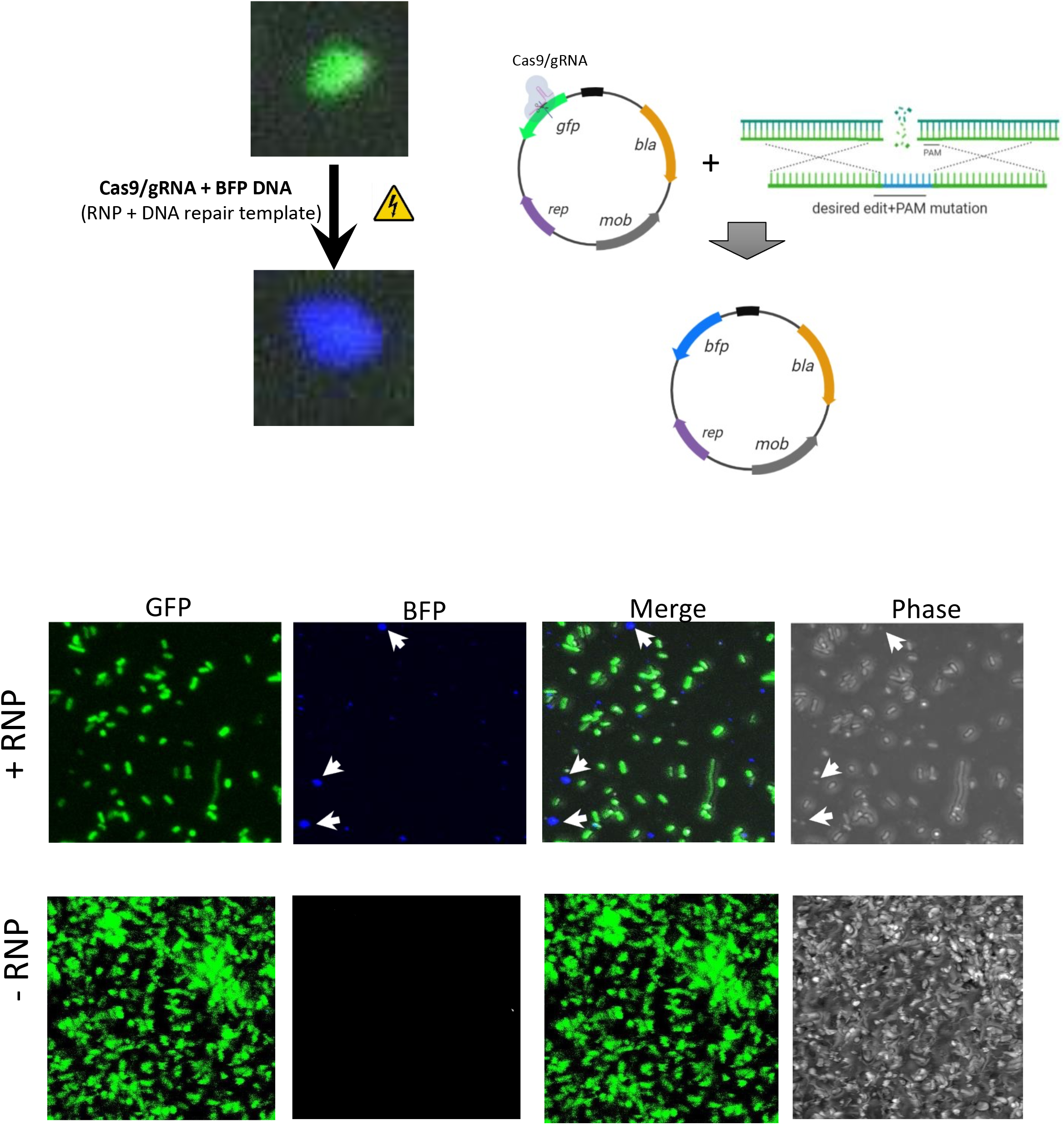
Cells with successful reparation produced fluorescence in color blue. Fluorescent cells can be observed after the electroporation (Fig. 5A). Although the observed efficiency was low (<5%), we could see a proof of principle for the use of RNPs in bacteria. As the transformation efficiency of SpCas9/gRNA complex would expectedly be limited due to its massive size, as we can see in the section B. Recombineering efficiency must be maximized.

## 5. Discussion

Since its discovery and application as a genome-modifying tool almost a decade ago [6], CRISPR-Cas has quickly become the standard gene-editing technology in eukaryotic and prokaryotic organisms. However, the versatility of CRISPR-Cas based tools in bacteria has not been as expanded as in eukaryotic organisms. One of these applications is the use of ribonucleoprotein sgRNA/Cas9 complexes and synthetic DNA oligonucleotide donors. The first requirement for the RNP method to work properly is a recombinant SpCas9 enzyme with precise and efficient DNA targeting and nuclease activity. Our 2-step purification protocol generated highly active SpCas9-6xHis from *E. coli* with a high enzymatic activity (95 %) to make a targeted double strand on a DNA substrate (Fig. 3). This result is superior to what others have reported for recombinant SpCas9 when purified by affinity chromatography alone [7], but equivalent to more laborious reports with three purification steps (IMAC, ion exchange and size exclusion chromatography) [8]. Our purification approach, therefore, enhances Cas9 specific nuclease activity without sacrificing protein yield. Moreover, the nuclease activity of our preparation was completely sgRNA dependent as SpCas9 alone did not trigger any cut on the DNA substrate (GFP gene).

In order to improve the observed low efficiency of in *vivo* gene-editing of Cas9/gRNA RNPs, despite a very high specific *in vitro* activity, several actions can be taken. Employing smaller Cas9 orthologs has been shown to radically improve the permeability of RNPs via electroporation [9]. Using bacterial strains expressing heterologous DNA recombination systems, e.g. Lambda Red, could also improve the efficiency of targeted gene-editing by the RNP method as has been shown for plasmid-based methods [10–12]. Another important aspect that can be improved is the RNP incorporation efficiency, especially in hard to electroporate cells such as thick-walled gram-positive bacteria. In this regard several approaches have been recently developed. These include the usage of cationic polymer-derivatized Cas9 [13] and lipid-Cas9 protein/mRNA nanoparticles [14,15].

The main advantage of the RNP approach is that it does not rely on the bacterial strain transcription and translation machinery [16], allowing to directly pre-evaluate the efficacy of the RNP preparation by *in vitro* nuclease assays as shown in the present work. Additionally, the RNP complex is usually degraded shortly after transfection, avoiding the toxic effects of a continuous Cas9 expression that has been described in several bacteria [17].

## Acknowledgements

We thank Enrique Alcaraz for technical assistance.

## Funding

EMR grants: SEP-PRODEP: CINVESTAV-PTC-002, “Ruben Lisker” Biomedicine grant 2017, CONACYT CB2017-2018: A1-S-10743, SEP-CINVESTAV 2018: 1.

RDAO: CONACYT Postdoctoral Scholarship: Application No. 2019-000006-01NACV-01211.

---

**Figure S1.**
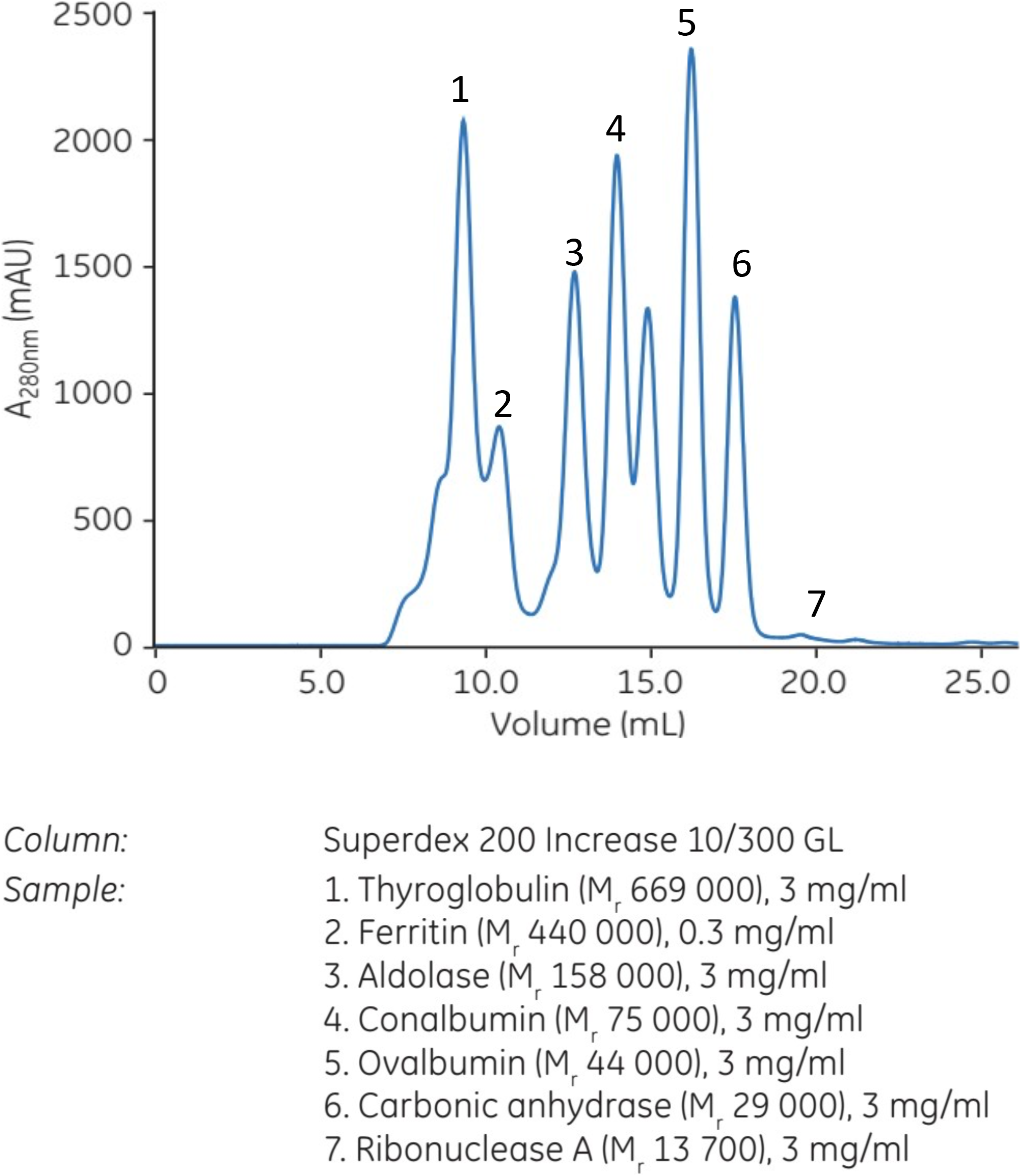
Gel filtration of molecular weight standards. Expected molecular size of several proteins on the Superdex 200 increase 10/300 GL column. Elution of SpCas9-6xhis (160 kDa) at 13.6 ml is consistent with its molecular weight according to this chromatogram, near to the elution time of aldolase, peak 3 with a MW of 158 kDa

## Notes

### Competing Interest Statement

The authors have declared no competing interest.

